# The Diameter Factor of Aligned Membranes Facilitates Cutaneous Wound Healing by Promoting Epithelialization in an Immune Way

**DOI:** 10.1101/2021.02.25.432929

**Authors:** Chenbing Wang, Chenyu Chu, Xiwen Zhao, Chen Hu, Li Liu, Jidong Li, Yili Qu, Yi Man

## Abstract

Topographical properties (such as pattern and diameter) of biomaterials play important roles in influencing cell activities and manipulating the related immune response during the skin wound healing. We prepared aligned electrospinning membranes with different fiber diameters including 319±100 nm (A300), 588±132 nm (A600) and 1048±130 nm (A1000). The A300 membranes significantly improved cell proliferation and spreading, and facilitated skin wound healing (epithelization and vascularization) with the regeneration of hair follicles. The transcriptome revealed the underlying molecular mechanism that A300 could promote macrophages transformation to reparative phenotypes with highly expressed IL10 and TGFβ, subsequently promoting epidermal cell migration via secreting matrix metalloproteinases 12 (MMP12). All the results indicate the A300 to be a potential candidate for guided skin regeneration applications.

**Graphic abstract:** 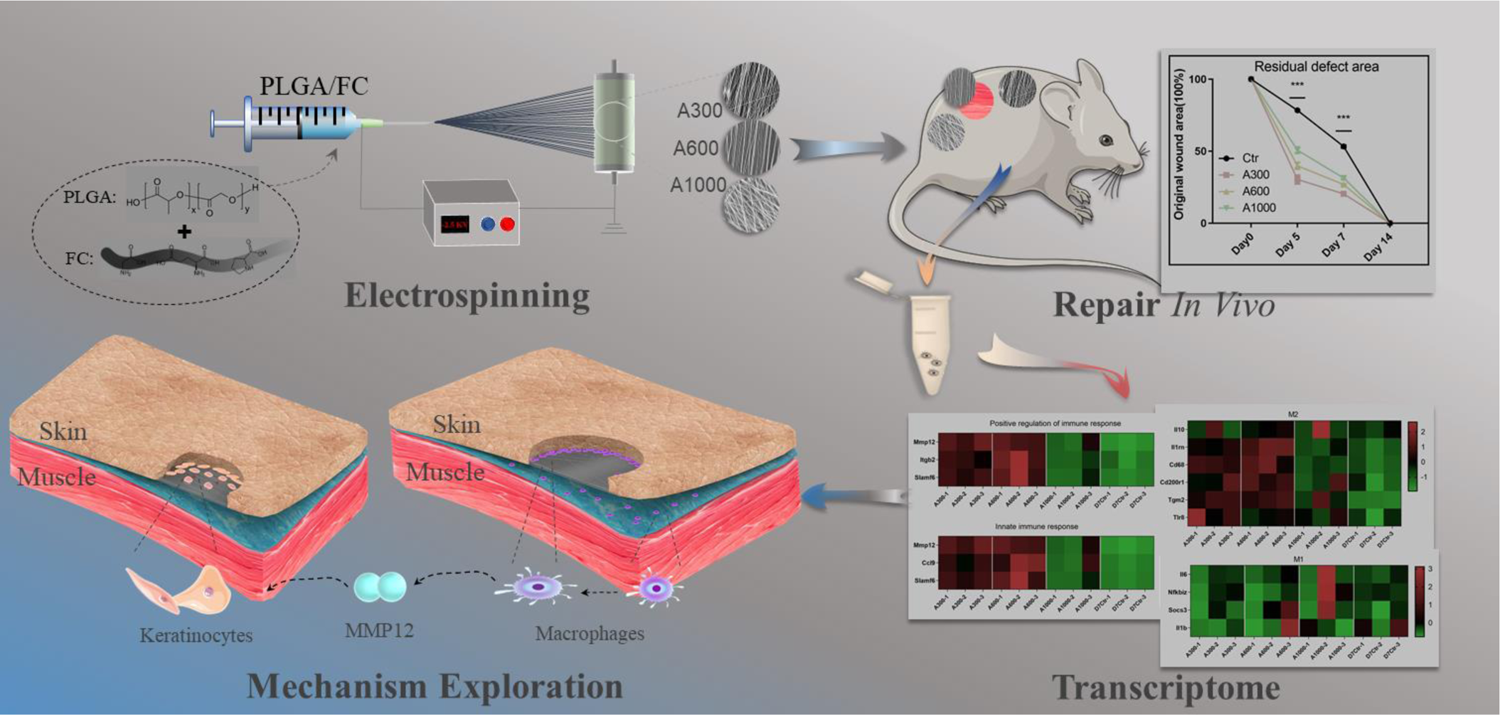

## 1. Introduction

Skin, as a superficial organ in contact with the surrounding environment, constitutes a first important guard-line against external hazards[1]. The inability to re-epithelialize wounded skin can contribute to dewatering, inflammation or even mortality[2, 3]. Therefore, it is very important to close the wound quickly to restore the barrier function essential for survival of the organism. Regrettably, current strategies are not satisfactory in managing large wounds, since they are dependent on both slow and passive healing processes[4]. Promoting skin healing with the regeneration of skin appendages such as hair follicles (HFs), that is closely related to skin tensile strength and one of the important indexes of skin functional healing[5], has not authentically been realized[6]. Consequently, there is a growing need for exploring advanced grafts to achieve ideal re-epithelialization with appendages[7–9].

Due to the characteristic of imitating the topography and functions of extracellular matrix (ECM)[10, 11], providing moist environment, allowing gas exchange, avoiding bacteria infiltration[12] and manipulating the polarization of macrophages towards an anti-inflammatory phenotype[13, 14], the electrospinning membrane is an ideal alternative for skin wound healing[15, 16]. Of these, the aligned membranes have a wide range of applications in the field of skin tissue engineering[1,17,18], since they can provide a series of biologically chemical and physical cues for regulating cell behaviors and influencing the immune response[19]. For example, studies have been reported that fibroblasts can migrate for a long distance in a highly correlated manner at a constant speed on the aligned membranes[20, 21]. Aligned membranes could promote the normal differentiation and outgrowth of vascular smooth muscle cells [17, 22]. Critically, the previous research in our group showed that aligned membranes have more advantages in soft tissue repair and can actively regulate immune response [23].

Even though already widely used as skin tissue engineering biomaterials, the aligned membranes are still verified with poor inflammation resistance and mechanical properties[24], which obviously impede its wider application. Many studies have attempted to improve physicochemical properties and biological performance of the aligned membranes via adjusting their microstructure, since topographical factors indeed exert some indispensable roles in early cell fate prior to cytokines[25, 26]. In view of the fact that the ECM is composed of different fibers from nano to micron, the diameter factor dictating the physicochemical properties and biological performances of membranes was introduced[27]. For example, human skin fibroblasts had a well-diffused morphology, growing on the membranes of the range of 350-1100 nm diameter, where the expression of type III collagen gene in human skin fibroblasts was significantly up-regulated[28]. It was found that there was a critical minimum diameter (d), namely 0.97 μm, making the human fibroblasts better directionally developing in contrast to d<0.97 μm[29]. Additionally, the membranes with the small diameter (about 250-300 nm) had stronger support for the dermal fibroblast proliferation than that with the diameter of about 1 μm[30]. Another study has been reported that the diameter of membranes could affect the immune response of macrophages, especially in the early stage of inflammation, since nanofiber membranes minimized inflammatory response relative to microfiber ones[31] and manipulated the macrophages towards a reparative phenotype[32].

Studies involving fibroblasts and macrophages have corroborated the positive effect of diameter factor of aligned membranes on cell behavior. However, the results of abovementioned researches are controversial. The specific characteristics of these aligned membranes as to which diameter interval is most suitable for skin tissue engineering need to be verified. And the potential diameter-mediated mechanism of repairing tissue defects has not been explored. Herein, this article innovatively explores the skin defect response to aligned bio-sythentic membranes with varying fiber diameters including 319±100 nm (A300), 588±132 nm (A600) and 1048±130 nm (A1000)[33, 34], to develop a more suitable surface wound healing medical device for manipulating the related immune response and promoting the re-epithelialization with appendages. This comprehensive study also evaluated the transcriptome of rat skin wounds on the aligned membranes of varying fiber diameters for exploring the potential diameter-mediated mechanism of repairing tissue defects.

## 2. Results and Discussion

### A300 improves the mechanical stability, hydrophilicity and degradation of aligned membranes

#### Topological and Mechanical Properties

Workflow for evaluating physicochemical properties of aligned membranes with different diameters was summarized in **Fig.1A**. The topology and corresponding fiber diameter distributions of membranes are shown as 319±100 nm (A300), 588±132 nm (A600) and 1048±130 nm (A1000) in **Fig. 1B** and **1C**, revealing that the fibers presented a homogeneously bead-less performance, and a highly aligned morphology (The electrospinning parameters were shown in **Table S1**. The preliminary experiments were shown in the **Fig. S1**). Based on the histogram of **Fig. 1C**, the mean diameter of fibers was reduced obviously with the change of operation and solution parameters. This is mainly because the increase of electric field promoted the stretch rate of electrospinning fibers and the decrease of poly (lactic-co-glycolic-acid) (PLGA) restrained the electrospinnability itself (**Table 1**), resulting in the electrospinning fibers with a narrower size distribution and smaller mean diameter[30]. It is typically considered that submicron-scaled bio-scaffolds possess better pro-healing effect for tissue engineering since the main advantage of submicron-scaled features over micron-scaled features is that they both provide a larger surface area to adsorb proteins and form more adhesion sites to integrin[35, 36]. The mechanical performance of aligned membranes is an important determinant for its application in wound healing, which is expected to present suitable mechanical strength during surgery and tissue regeneration. In this research, we evaluated the mechanical behaviors of different aligned membranes (before cross-linking) via tensile strength tests. The stress−strain curve (**Fig. 1D** and **1E**) illustrates that the tensile strength of small diameter aligned membrane (A300, 11.95±0.35 MPa) increased compared with medium diameter (A600, 6.80±0.49 MPa) and large diameter (A1000, 8.86±0.47 MPa). Meanwhile, the strain rate of A300 (64.73±3.51%) maintained a lower level (**Fig. 1F**) compared to A600 (101.3±2.37%) and A1000 (126.2±8.01%), apparently increasing the mechanical stability of aligned membranes, which could prevent scars in skin wound healing[37]. The mechanical performance of these membranes could be partially explained by the fracture process of fibers. This is primarily because membranes comprising smaller diameter fibers performed both higher strength and lower ductility[38, 39].

**Fig. 1.**
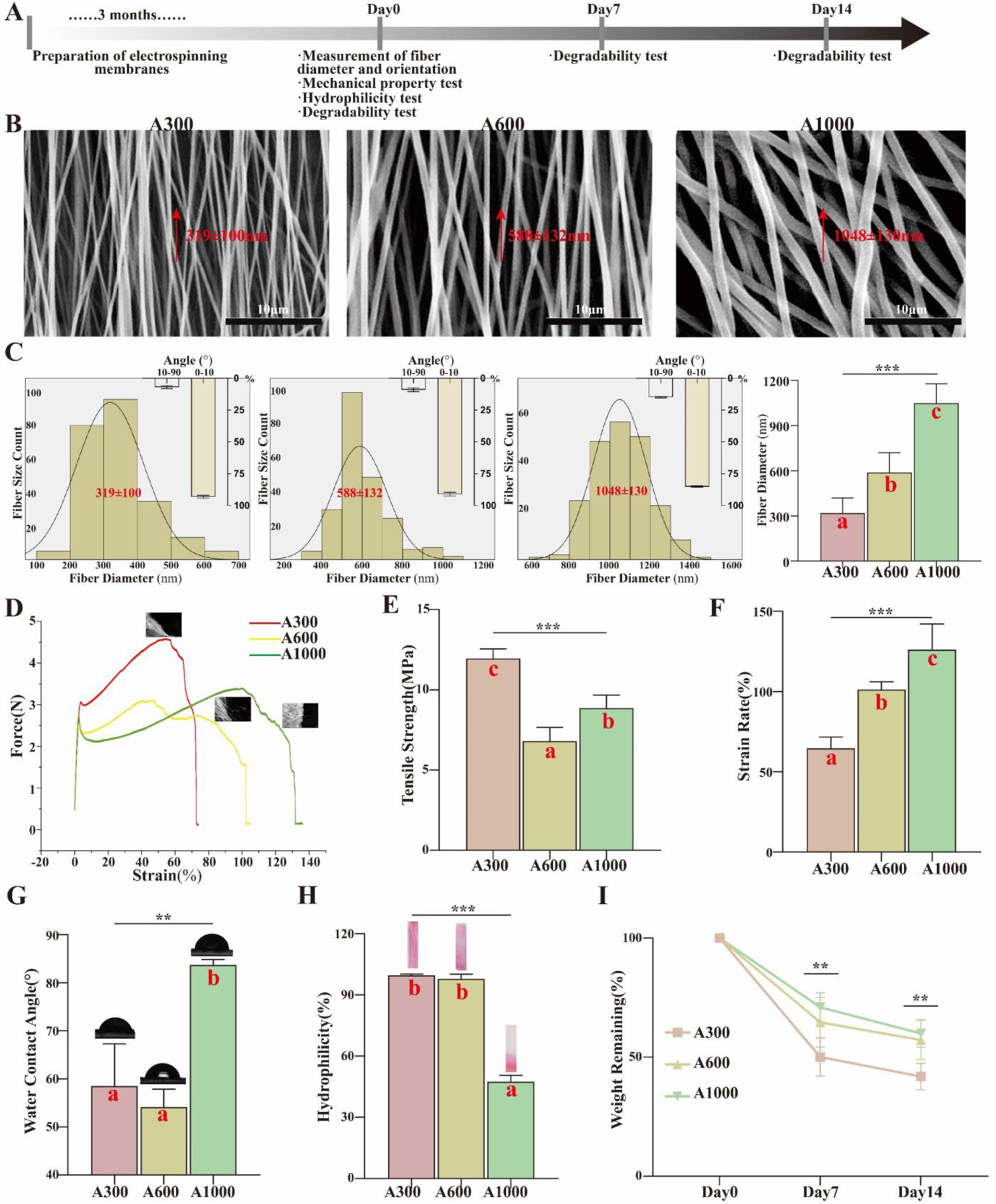
(A) Workflow for evaluating physicochemical properties of electrospinning membranes; (B) SEM images and (C) corresponding diameter distributions of different diameter aligned membranes; (D) Typical stress−strain curves of various membranes; (E) Tensile strength; (F) Strain rate; (G) WCA images and corresponding analysis; (H) Hydrophilicity images and corresponding analysis; (I) Degradation assays. Different letters indicate significant differences.

### Roughness and Physicochemical Properties

As shown in **Fig. S2A-C**, the roughness of aligned membranes seemed to be improved with the increase of mean diameter itself, consistent with the abovementioned result of SEM and beneficial to cell adhesion and migration to some extent[40]. Nevertheless, the result was not statistically significant difference. The aligned membranes possessed high specific surface area, thus providing larger surface area for cell adhesion[16], preventing undesirable fluid accumulation and accommodating more facile oxygen permeation[41]. It has been reported that membranes with fibers arranged in parallel (alignment) are beneficial to the normal differentiation of vascular smooth muscle cells, and have potential in application to the reconstruction of tissue-engineered blood vessels[17]. In this study, the alignment of A300 was similar to that of A600 yet much better than that of A1000, indicating its potential of vascularization (**Fig. S2D**).

### Hydrophilicity and *in Vitro* Degradation Behavior

The hydrophilicity of the aligned membranes was investigated by water contact angle (WCA) assay. As shown in **Fig. 1G**, the aligned membrane of A300 exhibited a water contact angle of 58.52°, resembling that of A600 (54.15°) yet less than that of A1000 (83.69°), which demonstrated its hydrophilic behavior. It has been proved that the decrease of fiber diameter significantly increased the water affinity of aligned membranes. This is mainly because the high specific surface area conveniently interacts with water molecules. The morphology of biomaterials directly dictated the hydrophilicity of biomaterials, ultimately determining the WCA. Apart from the fiber diameter, the fiber arrangement and surface roughness exerted significant influence on the hydrophilicity of aligned membranes. We further studied the hydrophilicity behavior by evaluating the infiltration of these three aligned membranes. As proved by **Fig. 1H**, the aligned membrane of A300 was completely wetted, marginally better than that of A600 (97.86%) yet significantly better than that of A1000 (47.54%), consistent with the aforementioned result of WCA.

Ideal implanted biomaterials such as guided tissue regeneration (GTR) membranes should have both suitable mechanical strength as well as matched biodegradability. Fiber degradation gradually destroys the structural and functional integrity of aligned membranes and provides the space and nutrients for new tissue ingrowth. In order to evaluating the degradability of the assayed membranes *in vitro*, we monitored the remaining mass during the immersion process. As shown in **Fig. 1I**, the remaining mass of A300 was significantly smaller than those of the others both on day 7 (A300/50.02±3.58%, A600/64.65±4.64% and A1000/70.97±2.73%) and day 14 (A300/41.84±2.47%, A600/57.24±3.71% and A1000/59.99±2.58%). The results showed that the decrease of diameter accelerated the degradation process of fibers, hence advancing the degradation of aligned membranes.

### 2.1. A300 promotes fibroblast proliferation and accelerates keratinocytes spreading on the aligned membranes

Workflow for evaluating the cell behavior of aligned membranes was summarized in **Fig.2A**. Before clinical application, the different diameter aligned membranes utilized for GTR should be considered to perform the ability for ameliorating related cell proliferation and/or migration. Accordingly, the proliferation of fibroblasts (L929) and human oral keratinocytes (HOK) on different diameter aligned membranes was assessed, as shown in **Fig. 2B** and **2C**. For L929, the proliferation rate of A300 was significantly higher than those of the others except for the control group all along. However, there was no statistically significant difference within groups of HOK proliferation. The results indicated that the small diameter membranes (A300) were more conducible for the proliferation of L929 yet seemed to have no significant influence on that of HOK [42]. It is reported that the cell performance on membranes are intimately related with fiber diameter and topological structure, directly manipulating the cell adhesion and exchange of nutrients[28,30,31]. The fiber diameter of the aligned membranes decreased from 1048±130 nm to 319±100 nm with the change of operation and solution parameters, promoting cell adhesion and proliferation of A300[43].

**Fig. 2.**
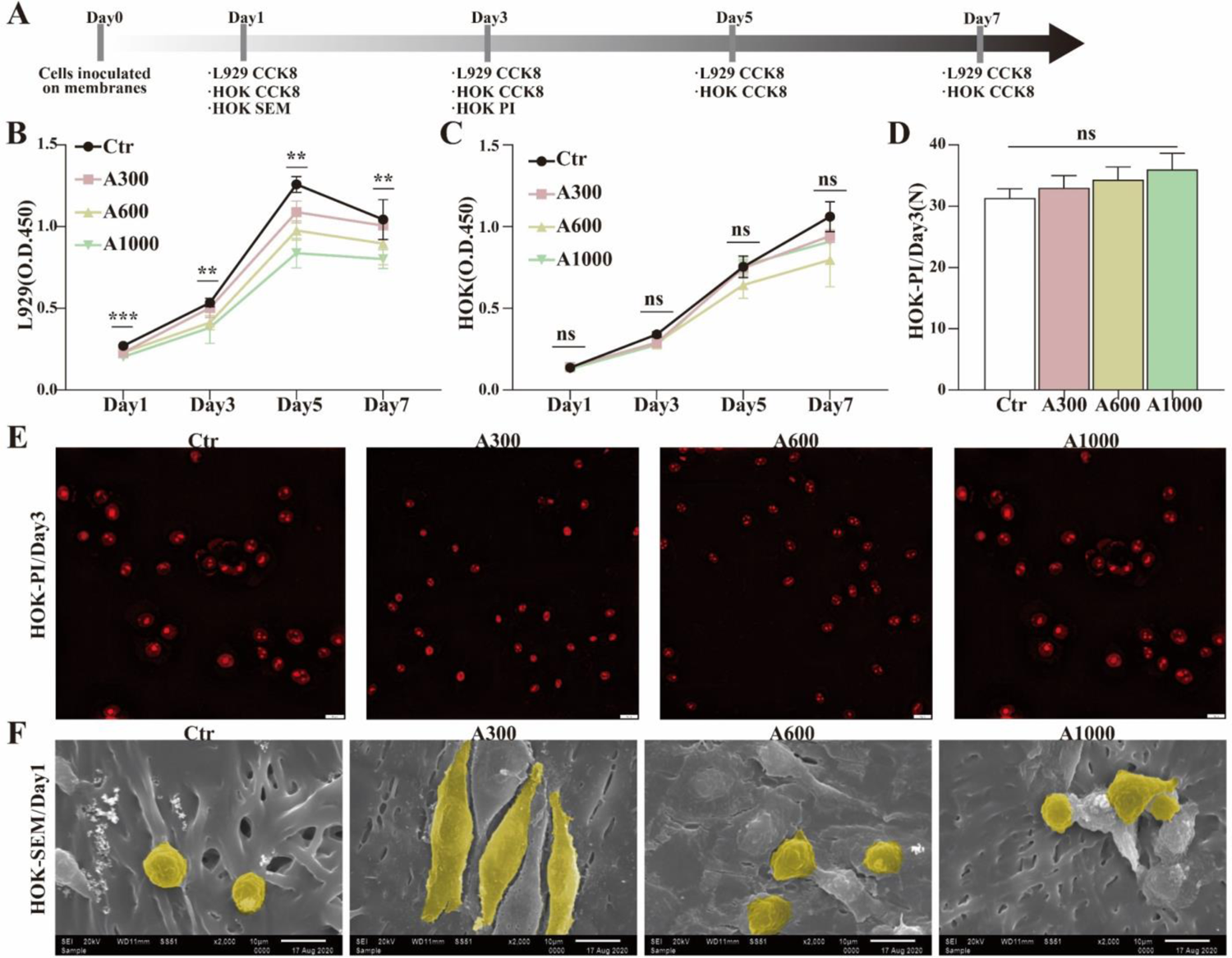
(A) Workflow for evaluating the biocompatibility of electrospinning membranes; CCK-8 tests of (B) L929 and (C) HOK; (D) Analysis and (E) corresponding images of PI fluorescent staining of HOK; (F) SEM images of HOK morphology.

### Viability and Morphology

In order to clarify the effect of aligned membranes with different diameters on HOK, we further studied the viability and morphology of HOK via the fluorescent staining of PI and SEM. As shown in **Fig. 2D** and **2E**, the dead HOK on aligned membranes of A300 (33.0±1.2) was marginally less than that of A600 (34.3±1.2) and A1000 (36.0±1.5) on day 3, without statistically significant difference. Additionally, as presented in **Fig. 2F**, the result of SEM revealed that the HOK spread on the aligned membranes of A300 much better than that of A600 and A1000 on day 1, consistent with previous studies that cells adhered on nanofibers performed much better than microfibers[44]. Overall, it may have different degrees of positive effects on different types of cells, such as promoting L929 proliferation and accelerating HOK spread, respectively. Our results also indicate that the surface properties of aligned membranes including diameter, wettability and roughness seemed to manipulate the biological behavior of L929 and HOK[45, 46].

### 2.2. A300 significantly facilitates skin wound healing

#### General Situation

Workflow for evaluating the rat skin wound healing and was summarized in **Fig.3A**. In practical application, the membranes utilized for GTR could be recognized to present the ability for accelerating related wound healing. Accordingly, the restoration performance of skin defect of rat on aligned membranes with different diameters was assessed, as shown in **Fig. 3B** and **3C**. For rat skin defects, the residual area of A300 group was significantly smaller than those of the others at all the time points. Intriguingly, the wounds in A300 group occasionally reached the complete healing one or two days earlier. The macroscopic result indicate that the small diameter membranes were indeed more conducible for the reconstruction of rat skin defects, completely consistent with the aforementioned results of cells *in vitro*. We further studied the wound healing by evaluating the hematoxylin and eosin (H&E) staining of tissue sections. As proved in **Fig. 3D** and **3E**, the residual gap of A300 group (2.70±0.18 mm) was significantly narrower than that of A600 (3.63±0.06 mm) and A1000 (4.15±0.05 mm) on day 7, confirming the previous macroscopic result that rat skin defects of A300 closed faster.

**Fig. 3.**
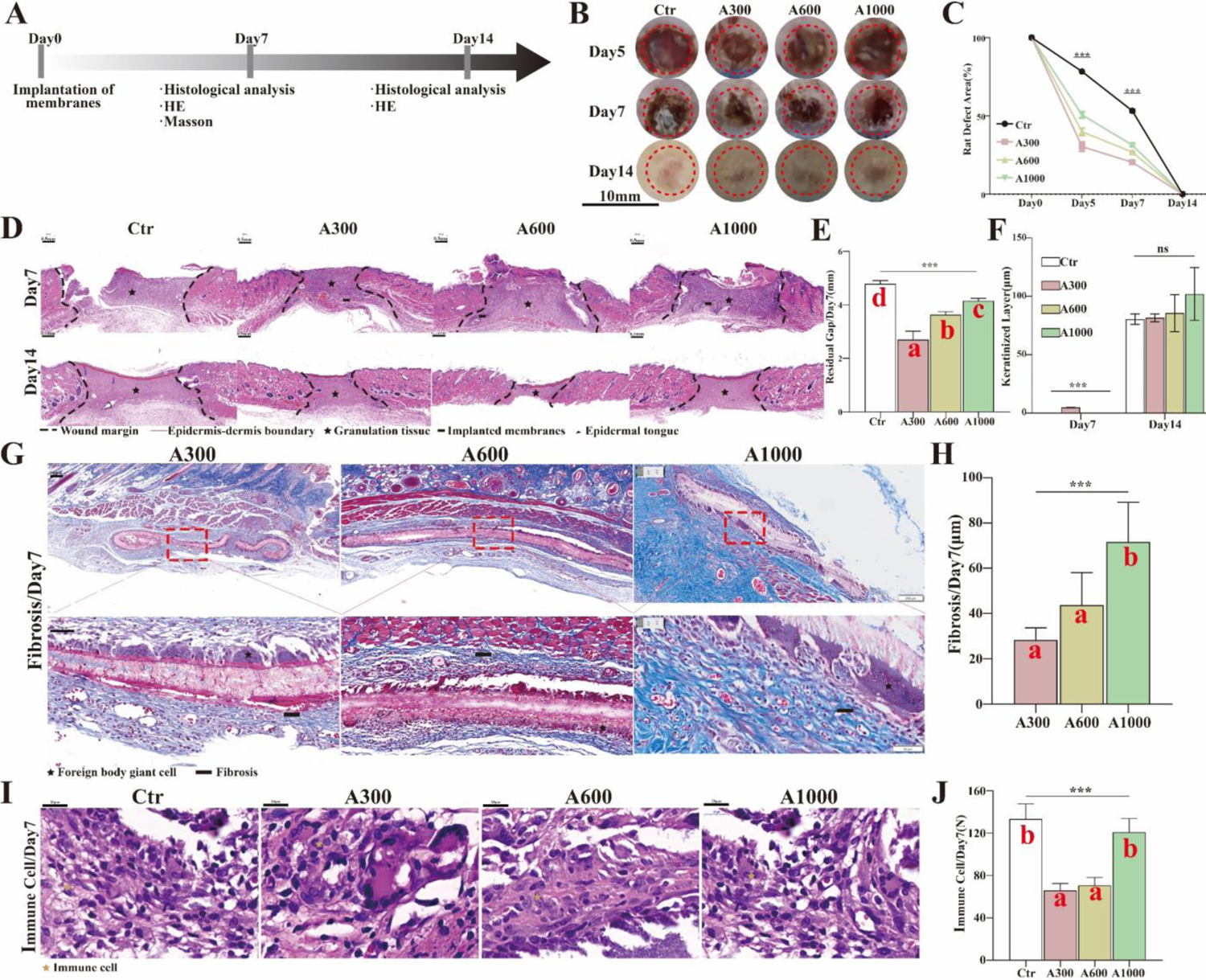
(A) Workflow for evaluating the rat skin wound healing and FBR; (B) Macroscopic images of rat skin defects and (C) corresponding analysis; (D) Residual gap images of H&E of rat skin defects and (E) corresponding analysis;(F) The analysis of keratinized layer images of H&E (G) FBR images of Masson staining of rat subcutaneous implantations and (H) corresponding analysis; (I) Inflammatory cell infiltration images of H&E of rat skin defects and (J) corresponding analysis. Different letters indicate significant differences.

#### Foreign body response (FBR)

The performance of aligned membranes is built on its compatible interaction with the host immune defense system[47], since the immune recognition initiates FBR comprising continuous inflammation, fibrosis (fibrous capsule) and damage to the surrounding tissue[48, 49]. These unwanted outcomes may destroy the function of aligned membranes and arise inevitable pain and discomfort on the recipient site[50]. In this study, the macroscopic result of rat skin defects showed that the better wound healing and less scar tissue formed in the A300 group on day 14 (**Fig. 3B**). We further researched the thickness of fibrous capsule of rat subcutaneous implantations via the MST staining to evaluate the degree of FBR. As depicted in **Fig. 3G** and 3**H**, the thickness of fibrous capsule of A300 group (28.40±2.38 μm) was significantly thinner than that of A600 (43.68±6.47 μm) and A1000 (71.66±7.83 μm) with statistically significant differences. Additionally, the level of immune cells recruited by aligned membranes of A300 (66.00±3.79) and A600 (70.67±4.33) was less than that of A1000 (121.0±7.37) (**Fig. 3I** and **3J**). It’s ultimately concluded that aligned membranes of A300 could compatibly interact with the host immune defense system and marginally give rise to the FBR for reaching the normal healing. Other researchers have described that nanofiber scaffolds minimized inflammatory response relative to microfiber scaffolds[31].

#### Vascularization

It’s well acknowledged that the newly formed blood vessels could provide the necessary nutrients utilized for promoting wound healing[51]. Workflow for evaluating the vascularization and epithelization was summarized in **Fig.4A**. Based on images of **Fig. 4B** and **4C**, the results indicated that the degree of vascularization of A300 group (0.250±0.009 mm^2^) was significantly greater than that of A600 (0.125±0.002 mm^2^) and A1000 (0.080±0.003 mm^2^) on day 7, as further demonstrated by the dominant IHC staining (**Fig. 4D** and **4E**) and corresponding RNA expression of CD31 in A300 group (**Fig. 4F**).

**Fig. 4.**
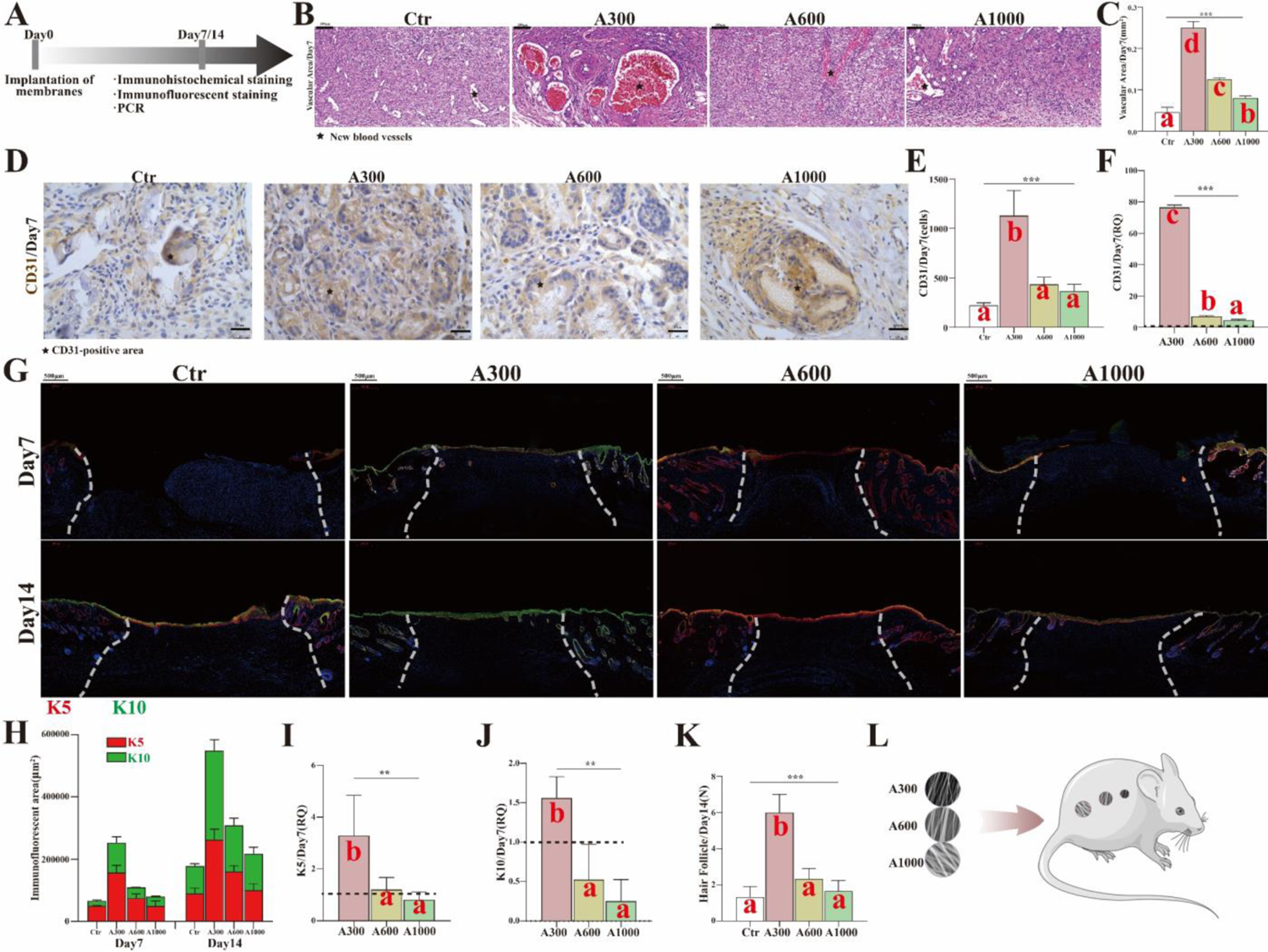
(A) Workflow for evaluating the vascularization and epithelization; (B) Vascularization images of H&E of rat skin defects and (C) corresponding analysis; (D) CD31-positive images of IHC staining of rat skin defects and (E) corresponding analysis; (F) The RNA expression of CD31; (G) K5(red)/K10(green)-positive images of IF staining of rat skin defects and (H) corresponding analysis; (I) The RNA expression of K5; (J) The RNA expression of K10; (K) The number of regenerated hair follicles on day 14; (L) Different diameter aligned electrospinning membranes implanted on rat skin defects. Different letters indicate significant differences.

#### Epithelization

Additionally, the keratinized layer of wounds first appeared in A300 group (4.78±0.08 μm) yet didn’t exist in other groups on the 7th day after operation. There was no significance between control and various diameter groups on day 14, whereas the thickness of keratinized layer in A300 group was closer to that in control group (**Fig. 3F**), as further corroborated by the IF staining (**Fig. 4G** and **4H**) and corresponding RNA expression of K5/K10 (**Fig. 4I** and **4J**), both revealing that the degree of epithelization (continuity and differentiation) of A300 group was significantly better than other groups. Hair follicle regeneration is one of the important indexes of skin functional healing[5]. It has been reported that hair follicle stem cells could accelerate wound healing as well as tensile strength[52]. And the maximum number of newly-formed hair follicles on day 14 (**Fig. 4K**) was presented in A300 group, demonstrating that A300 group achieved excellent repair results with regenerated appendages[53]. It is so concluded that the A300 group played a more important role in both vascularization and epithelization, suggesting that aligned membranes of A300 indeed accelerated wound healing of rats. However, the specific mechanism of reducing inflammation/FBR and subsequently promoting skin wound healing remains unclear.

### 2.3. MMP12 upregulated by A300 promotes keratinocytes migration

In order to further explore the specific mechanism of reducing inflammation/FBR for the better wound healing, the Bulk RNA sequencing of rat skin defects implanted with different diameter aligned membranes was studied via the principal component analysis (PCA), Gene Ontology (GO) and Kyoto Encyclopedia of Genes and Genomes (KEGG) to screen the inflammation-related targets. Workflow for exploring the specific mechanism was summarized in **Fig.5A**. The results of PCA showed that A300 was obviously different from other diameter and control groups (**Fig. 5B**), suggesting that the diameter factor indeed played a vital role in wound healing. As presented in **Fig. 5C-E**, the results of GO at the front showed that the significantly up-regulated biological process (BP) items of A300 primarily included immune-related processes, such as *positive regulation of immune response* and *innate immune response*, relative to control and A1000 groups. However, compared with A600, the significantly up-regulated BP items of A300 was angiogenesis- and extracellular matrix-related process, which probably resulted from the less difference of wound healing between A300 and A600.

**Fig. 5.**
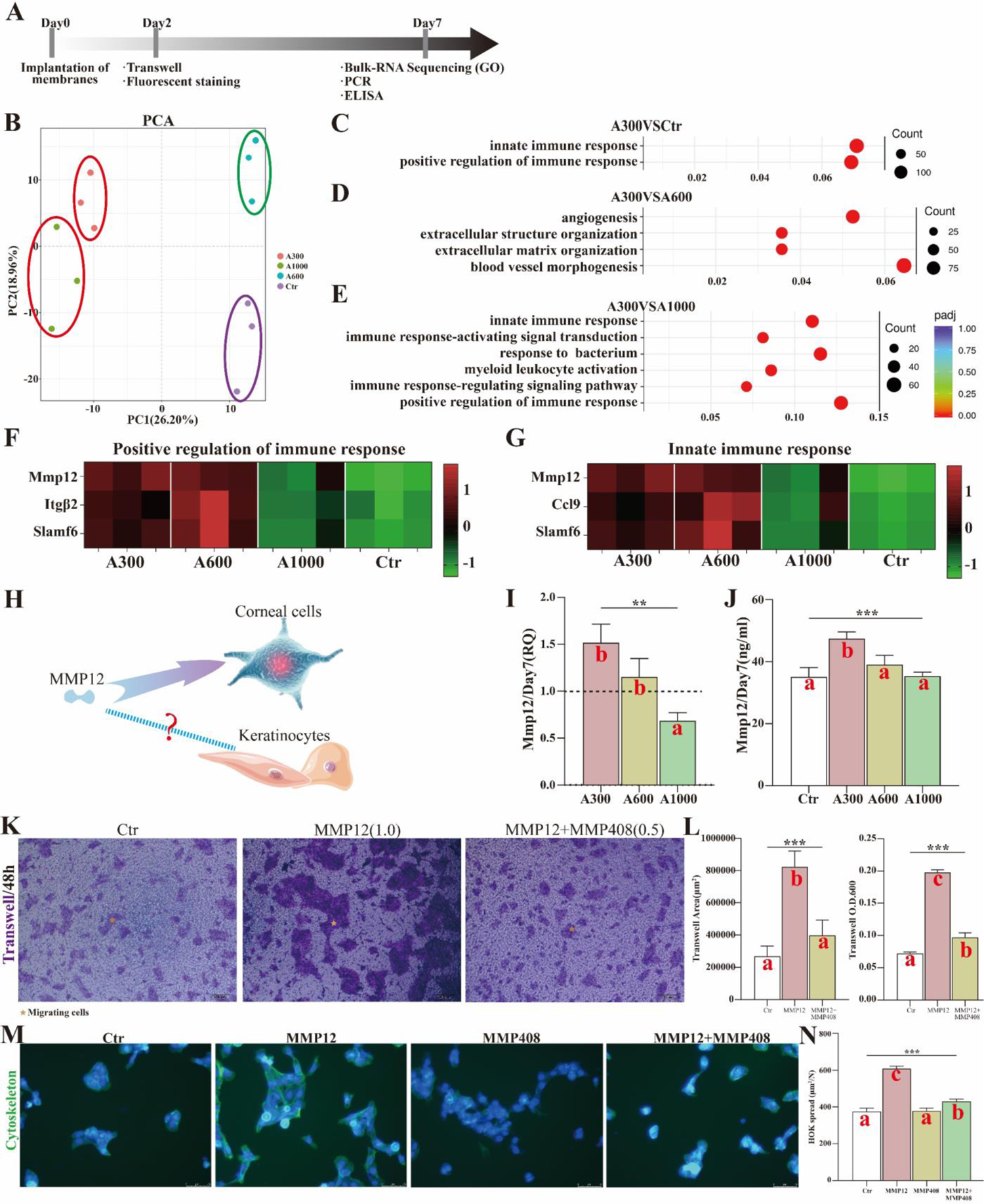
(A) Workflow for exploring the function of MMP12 during the rat skin defects; (B) PCA images of Bulk RNA Sequencing on day 7; (C) A300*VS*Ctr, (D) A300*VS*A600 and (E) A300*VS*A1000 of GO of Bulk RNA Sequencing;(F) Heat map of *positive regulation of immune response*; (G) Heat map of *innate immune response*; (H) The speculation of MMP12 function; (I) The RNA expression of MMP12; (J) ELISA result of MMP12; (K)The trans-well assays on MMP12 promoting the migration of HOK and MMP408 inhibiting the function of MMP12 and (L) corresponding analysis; (M)The fluorescent staining on MMP12 promoting the spread of HOK and MMP408 inhibiting the function of MMP12 and (N) corresponding analysis. Different letters indicate significant differences.

After that, the genes of immune-related processes that displayed significant changes were chosen and presented as heat maps. From the heat maps (**Fig. 5F and 5G**), we learned that the expression of A300 was similar to that of A600 yet much stronger than that of control and A1000 in the abovementioned immune-related process, indicating that A300 could reduce inflammation/FBR probably by actively regulating immune response for achieving better wound healing. More importantly, the significantly differentially expressed matrix metalloproteinases 12 (MMP12) appeared both in *positive regulation of immune response* and *innate immune response* of rat skin defects. It has been reported that the tissue inhibitor of matrix metalloproteinase (TIMP)-1 significantly reduced the migration ability of keratinocytes[54]. Another study has revealed that MMP12 actively regulated the migration of cells and protects against corneal fibrosis in the process of corneal wound healing[55, 56] (**Fig. 5H**). We wondered whether MMP12 had a similar role in skin wound healing. For this reason, we evaluated the RNA and protein expression of MMP12 of rat skin defects implanted with different diameter aligned membranes through qRT-PCR and ELISA. The RNA and protein expression of MMP12 of A300 were higher than other groups and reflected statistically significant difference (**Fig. 5I** and **5J**). From the results of Tran-swell, there were more migrating and more spread HOK in MMP12 group than in MMP12 plus MMP408 (the inhibitor of MMP12) and control group, which revealed that the recombinant protein MMP12 (The screening of the optimal concentration of MMP12 for HOK migration was shown in **Fig.S3**) promoted the migration and spread of HOK and the MMP408 inhibited the promoting migration and spread effect of MMP12 (**Fig. 5K-N**), indicating that MMP12 indeed facilitated the migration and spread of epidermal cells.

#### A300 promotes macrophages transforming to reparative phenotypes

Workflow for exploring the transformation of macrophages was summarized in Fig.6A. It’s commonly considered that MMP12 is predominantly expressed by macrophages[57] (**Fig. 6B**). The phagocytosis, as a process for maintaining cell homeostasis and organelle renewal, is a pivotal factor in inducing macrophages to reprogram from pro-inflammatory phenotype (M1) to pro-healing and pro-resolving phenotype (M2) during the wound healing, as alternatively activated pathways play little role in M1-M2 reprogramming in vivo[58, 59]. As for this research presented in **Fig. 6C-E**, the results of KEGG at the front showed that the significantly up-regulated signal pathways items of A300 primarily included *lysosome* and *phagosome*, relative to control and A1000 groups. The heat maps derived from *lysosome* and *phagocytosis* items (**Fig. 6F** and **6G**) also presented genes related to phenotype transformation of macrophages. Additionally, the macrophage-related genes that displayed significant changes were chosen and presented as heat maps. We acquired that the expression of A300 on reparative macrophages (M2) was similar to that of A600 yet much stronger than that of control and A1000 according to heat maps (**Fig. 6H**). It has been reported that M1 could inhibit keratinocyte migration, harmful to wound healing[54, 60].The results of RNA expression on Cd68, IL10 and TGFβ (**Fig. 6I**) also showed that A300 was much more pronounced than other groups, corroborating that aligned membranes of A300 indeed promoted macrophages transforming to M2 (**Fig. S4**). Moreover, the autophagy marker, LC3B (IHC staining) of rat skin defects (**Fig. 6J**-**K**) corroborated the above results of gene heat maps. All the results indicated that A300 promoted macrophages transforming to M2.

**Fig. 6.**
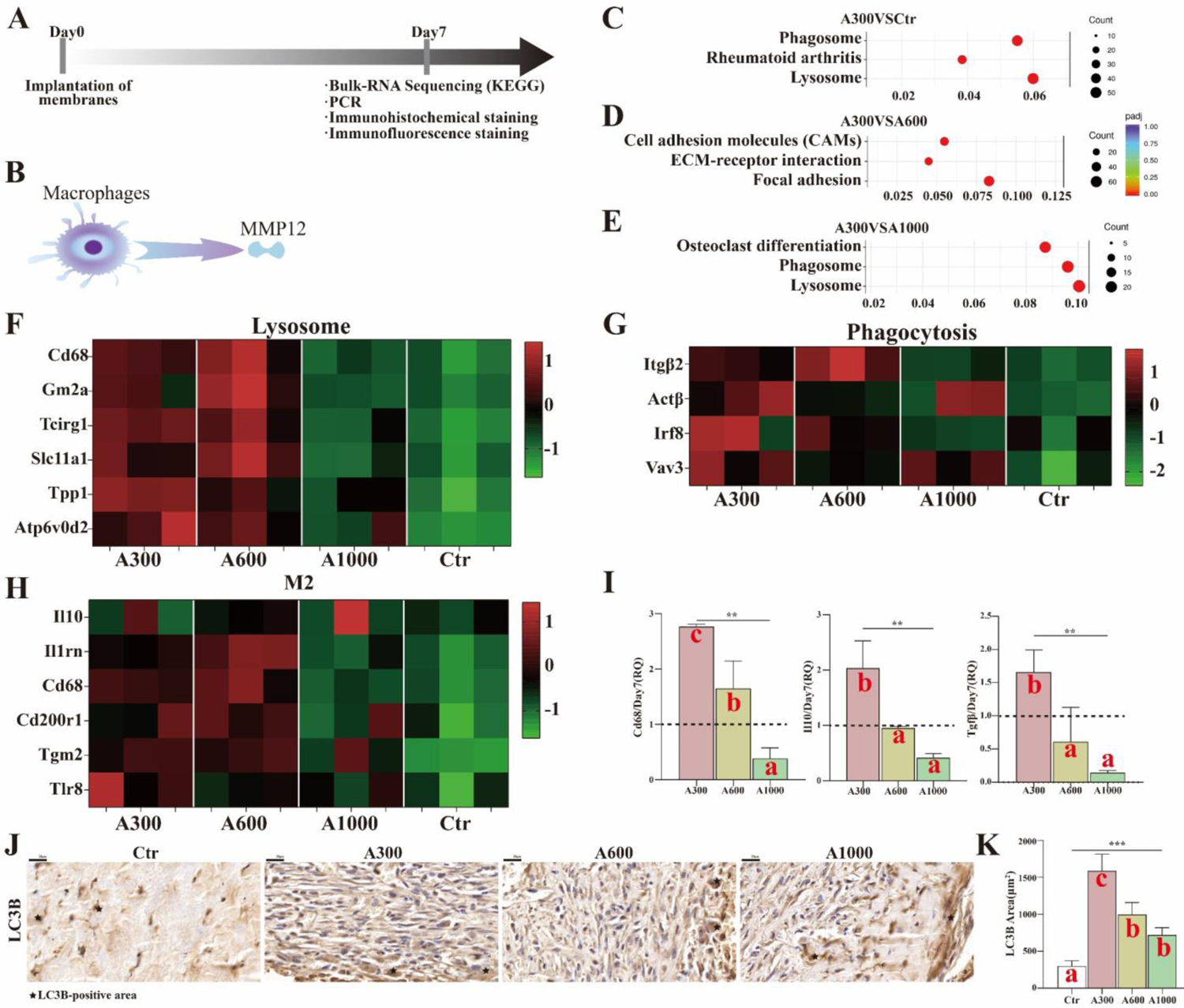
(A) Workflow for exploring the transformation of macrophages; (B) The MMP12 secreted by macrophages; (C)A300*VS*Ctr, (D) A300*VS*A600 and (E)A300*VS*A1000 of KEGG of Bulk RNA Sequencing on day 7; (F) Heat map of lysosome; (G) Heat map of phagocytosis; (H) Heat map of M2; (I) The RNA expression of Cd68, IL10 and TGFβ; (J) LC3B-positive images of IHC staining of rat skin defects and (K) corresponding analysis. Different letters indicate significant differences.

### 2.4. MMP12 secreted by macrophages on the A300 promotes keratinocytes migration

We further explored the effect of electrospinning membrane materials on macrophages. Workflow for verifying the function of macrophages and MMP12 *in vitro* was summarized in **Fig.7A**. The results of IF staining showed that the adhesion and extension of macrophages (Area/Number) on A300 group were better than other groups (**Fig. 7B** and **7C**), suggesting that A300 manipulated the behavior of macrophages. Through the cell scratch test, the macrophage conditioned medium of A300 group promoted the migration of keratinocytes (**Fig. 7D** and **7E**). Furthermore, the expression of MMP12 and M2 marker of macrophages was more significant in A300 group (**Fig. 7F-H**). More importantly, the recombinant protein MMP12 could enhance the promoting migration effect of macrophage conditioned medium of A300, while MMP12 inhibitor could largely inhibit this effect (**Fig. 7I** and **7J**). All the results indicated that MMP12 secreted by macrophages promoted keratinocytes migration.

**Fig. 7.**
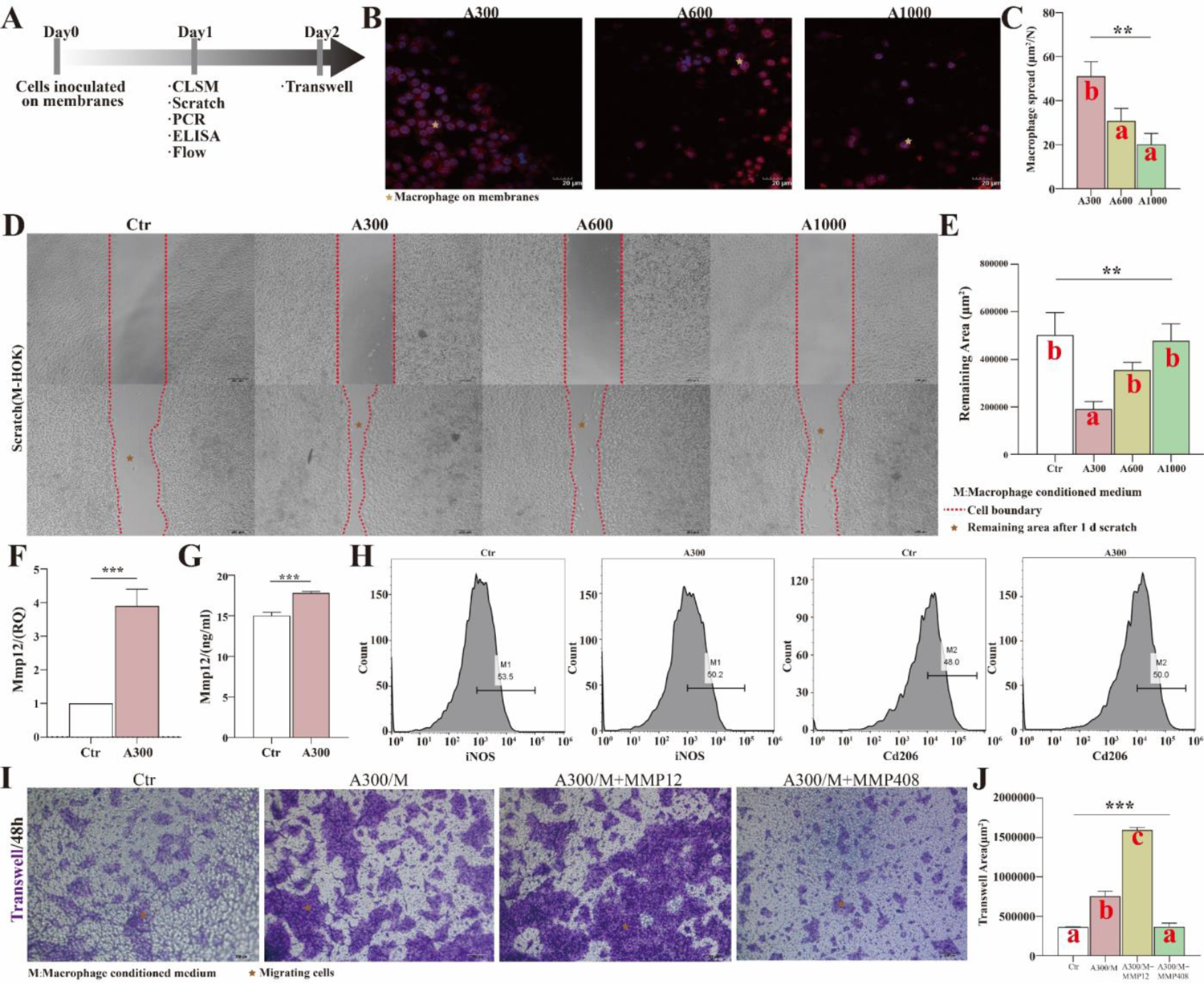
(A) Workflow for verifying the function of macrophages and MMP12 *in vitro*; (B) The CLSM images of macrophages (Thp1) on different diameter electrospinning membranes and (C) corresponding analysis; (D) The migration of HOK was stimulated by the conditioned medium derived from macrophages inoculated on various membranes and (E) corresponding analysis; (F) The RNA expression of MMP12; (G) ELISA result of MMP12; (H) The flow results of macrophages; (I) The trans-well assays and (J) corresponding analysis. Different letters indicate significant differences.

Workflow for verifying the function of MMP12 in mice was summarized in **Fig.S5**, showing that proper concentrations of MMP12 could accelerate epithelialization. It is so inferred that aligned membranes of A300 group could promote macrophages transformation to M2 through phagocytosis, subsequently promoting epidermal cell migration and wound healing by secreting IL10, TGFβ and MMP12.

## 3. Conclusions

In present studies, the aligned membranes with different diameters were successfully prepared via electrospinning, which is capable to imitate the skin component and spatial structure of the ECM. The 319±100 nm (A300) group significantly enhanced the mechanical strength and hydrophilicity, and accelerated the degradation rate of the fabricated aligned membrane as desired. The A300 promoted L929 proliferation and accelerated HOK spreading, respectively; in particular, the A300 facilitated skin wound healing (epithelization and vascularization). Above all, the transcriptome analysis revealed the underlying molecular mechanism that aligned membranes of A300 group could promote macrophages transformation to M2 through phagocytosis, subsequently exerting anti-inflammation via secreting IL10, TGFβ and promoting epidermal cell migration via secreting MMP12. Hence, we consider the small diameter aligned fibrous membrane (A300) to be a potential candidate for guided skin regeneration applications, particularly for the epithelization.

## 4. Experimental section

### 4.1. Materials

Poly (lactide-co-glycolide) (PLGA, LA/GA = 75:25, Mw = 105 kDa, dispersity is 1.897) was purchased from Jinan Daigang Biomaterial Co., Ltd. (Shandong, China). Fish collagen (FC) from fish scale and skin was procured from Sangon Biotech Co., Ltd. (Shanghai, China). 1,1,1,3,3,3-Hexafluoro-2-propanol (HFIP), N-hydroxysuccinimide (NHS), and 1-ethyl-3-(3-dimehylaminopropyl) carbodiimide hydrochloride (EDC) were offered by Aladdin Co., Ltd. (Shanghai, China).

### 4.2. Preparation of aligned membranes with different diameters via electrospinning

To fabricate the aligned fish collagen reinforced Poly (lactide-co-glycolide) electrospinning membrane (FC/PLGA) with different diameters, PLGA and FC were dissolved in HFIP under stirring at 25°C. The solution was stirred vigorously until complete dissolution of PLGA and FC. Subsequently, the prepared electrospinning solutions were employed to fabricate fibrous aligned membranes with different diameters via electrospinning. The electrospinning solutions were loaded into a plastic syringe fitted with a flat-tipped 21 G stainless steel needle. A high voltage and a distance were applied between the needle and the roller collector of 2800 rpm covered with a piece of aluminum foil. The solution was fed at a constant speed, regulated by a precision syringe pump. The prepared electrospinning membranes were dried in a vacuum oven at 25°C until the solvent was completely volatilized, which was confirmed by the X-ray diffraction (XRD) and fourier transform infrared spectroscopy (FTIR) spectra (**Fig. S2E and S2F**).

### 4.3. Characterization

Scanning electron microscope (SEM; JEOL, JSM-6510LV, Japan) was employed to observe the morphology of the aligned membranes with different diameters. In addition, Image-Pro Plus was applied to quantitatively measure the randomly selective 200 fiber diameter, distribution and alignment (orientation<10°)[61], from the SEM images obtained.

To estimate the mechanical performance of these membranes, the samples were attached to an electronic universal testing machine (SHIMADZU, AG-IC 50 KN, Japan) with a 50-N load cell. Prior to testing, the samples with thickness of 0.1−0.2 mm were cut into a dumbbell shape having a gauge length of 75 mm and width of 4 mm. The tensile properties of membranes were determined under a constant upper clamp speed of 5 mm/min at room temperature in accordance with the criteria of‘Plastics-Determination of tensile properties of films’(GB/T 1040.3−2006, corresponding with ISO 1184−1983). The elastic modulus was calculated from the slope of the linear region (ε = 1−3%) of the tensile-stress curve. Atomic force microscope (AFM; JEOL, JSM-6510LV, Japan) was employed to observe the roughness of the aligned membranes with different diameters. The surface wetting behavior of the membranes was characterized by measuring the water contact angles by using a contact angle measuring instrument (Chengde Dingsheng, JY-82B, China) and hydrophilicity test. Five samples were tested for each type of membrane to obtain an average value.

### 4.4. *In vitro* swelling and degradability study

The aligned membranes with different diameters were immersed in 50 mM of EDC/NHS and 10 mM of MES ethanol solution for 24 h at 4°C. The membranes were washed three times with ethanol and then dried in vacuum oven for 24 h. The dried fibrous membranes were cut into squares of side 10 mm and weighed accurately; then, they were put into a 5 mL plastic tube with 4mL of phosphate buffer solutions (PBS, pH = 7.4). Subsequently, all tubes were put in a shaking incubator at 100 rpm and 37°C. The remaining incubating media were changed every week. At each predetermined time point, the samples were dried to a constant weight in a vacuum oven. The swelling and weight loss rate were calculated according to the following formula:

Weight remaining (%) = m / m0×100%

Where m_0_ is the mass of membranes before incubation and m is the mass of membranes after incubation for different time.

### 4.5. *In vitro* cell culture experiments

#### 4.5.1. Cell culture

The fibroblasts (L929), human oral keratinocytes (HOK) and Thp1 were obtained from West China School of Stomatology, Sichuan University. The extraction was according to the normative procedure, which was approved by the Ethics Committee of Sichuan University. The L929, HOK and Thp1 were cultured in RPMI medium modified (1640; Gibco, USA) supplemented with 10% fetal bovine serum (FBS, Gibco, USA) and a 1% mixture of penicillin/streptomycin (MP Biomedicals, USA). The culture medium was replaced every other day in the L929, HOK and Thp1 culture cycle. On reaching a confluence of 80% to 90%, the Thp1 was induced into macrophages with the propylene glycol methyl ether acetate (PMA, 100 ng/ml). These membranes were preprocessed by immersing in a 75% ethanol solution for 0.5 h. Subsequently, all the prepared membranes were sterilized by γ-irradiation for cell experiment.

#### 4.5.2. Cell viability

The L929, HOK and macrophages (1×10^4^ cells/well, respectively) were seeded and incubated separately at 37°C with 5% CO^2^ atmosphere. One day after culture on membranes, the macrophage supernatant was collected for follow-up experiments. At the given time points (1 d, 3 d, 5 d, 7 d), the proliferation of L929 and HOK on these membranes was assessed by cell counting kit-8 (CCK-8, DOJINDO, Japan) assay (Membrane materials were not used in the control group). The optical density (OD) value was read by a 1420 Multilabel Counter (PerkinElmer, USA) at 450 nm. The dead portion of HOK was stained with propidium iodide (PI), according to the product instructions (DOJINDO, Japan). Fluorescence images were visualized with an inverted fluorescence microscope (Leica, Germany). Scanning electron microscopy (SEM; JEOL, JSM-6510LV, Japan) was employed to observe the morphology of cell on the aligned membranes (Random membrane materials were used in the control group). The macrophages were stained with phalloidin, according to the product instructions (Solarbio, China). Fluorescence images were visualized with the confocal laser scanning microscope (CLSM, Leica, Germany).

#### 4.5.3. Cell migration

The HOK (1×10^5^ cells/well) were seeded in the chamber of Trans-well and 24-pore plate at 37°C with 5% CO^2^ atmosphere. Different stimuli, such as the recombinant human matrix metalloproteinases 12 (MMP12, 1 μl, 20 ng/ml, Biolegend, U.S.A), MMP408 (the inhibitor of MMP12, 1 μl, 20 ng/ml, Invivogen, U.S.A) and macrophage supernatants were added. At the given time points (2 d), the HOK in the chamber of Trans-well and 24-pore plate were fixed (30 min). The HOK in the chamber of Trans-well stained with crystal violet (w/v0.1%, 30 min) and washed with PBS, finally visualized with the inverted microscope (Leica, Germany). The HOK in the 24-pore plate stained with phalloidin, according to the product instructions (Solarbio, China). Fluorescence images were visualized with the confocal laser scanning microscope (CLSM, Leica, Germany).

#### 4.5.4. Flow cytometry

Macrophages (transformed from Thp1) were collected into the EP tube after 24 h, fixed with 4% paraformaldehyde for 15 min, centrifuged and washed with PBS for 3 times; blocked with 3% BSA. Macrophages were mixed with the corresponding direct labeled antibody (iNOS, ab115819, 1:200; Cd206, ab195192, 1:200) prepared with 1% BSA solution, incubated at 4℃ for 1 h and washed with PBS for 3 times. The corresponding indicators were detected on the flow cytometer (Thermo fisher, U.S.A).

### 4.6. Animal experiments

Sprague−Dawley (SD) rats aged 8 weeks with an average weight of 220±20 g were obtained from the experimental animal center of Sichuan University approved by Institution Review Board of West China Hospital of Stomatology (No. WCHSIRB-D-2017-033-R1). The experimental rats could touch feed and tap water at will. The healthy status of the rats was checked every day during the experimental period. After conventional fasting for 12 h, the experimental rats were intraperitoneally anaesthetized in accordance with the instructions of the Animal House. Subsequently, a circular defect of diameter 6mm was created on the right and left skin of the dorsum via surgery following skin preparation. After transplanting different diameter aligned electrospinning membranes, the skin wounds were covered with 3M™Tegaderm™ and fixed on the latex ring with 3.0 silk suture. The experimental rats should receive routine postoperative nursing[62]. The experimental rats were divided into four groups with randomness: skin wounds transplanted with the saline (Ctr), A300 aligned membrane, A600 aligned membranes and A1000 aligned membranes. The rats were executed by dislocation after anesthesia on 5, 7 and 14 days after surgery to harvest the tissue samples for general analysis. All tissue sections were obtained from the rat skin wounds for routine histological examination and stained with Masson’s trichrome staining (MST Staining). And the thickness of fibrous capsule was determined manually under a microscope, using 5 randomly picked fields (at 100 magnification) of the Masson’s sections for evaluating the FBR. Immunohistochemistry (IHC) was performed for CD31 (ab182981, Abcam, 1:2000) to evaluate the vascularization, and LC3B (Sigma-Aldrich, L7543, 1:200, U.S.A) to evaluate the autophagy. Double staining immunofluorescence (IF) was performed for cytokeratin 5 (K5, ab52635, Abcam, 1:200) and cytokeratin 10 (K10, ab76318, Abcam, 1:150) to evaluate the keratinization. Both were further verified by quantitative reverse transcription-polymerase chain reaction (qRT-PCR), of which the data were presented as the relative quantification (RQ<2^-ΔΔCt^) compared with control groups (The related primers were shown in **Table S2**).

### 4.7. Bulk RNA sequencing

Three replicates of rat skin wounds in each group were detected for the assay of Bulk RNA sequencing (NEBNext® Ultra™ RNA Library Prep Kit for Illumina®), and the results were analyzed for Gene Ontology (GO) and Kyoto Encyclopedia of Genes and Genomes (KEGG) for screening the related targets. These were further verified by fluorescence quantitative PCR (The related primers were shown in **Table S2**) and enzyme linked immunosorbent assay (ELISA, MEIMIAN, China).

### 4.8. Statistical analysis

At least three independent assays were conducted to obtain repeatable data if not distinctively explained, and the data was dealt with Case Viewer, Origin 2019 or Graphpad Prism 8.0 software. Statistical analyses were performed by Student’s t-test and the Tukey post-hoc test by analysis of variance under the condition of normal distribution. The numerical data are presented as mean ± standard error of the mean. A value of p < 0.05 was considered statistically significant (\**p* < 0.05, \*\**p* < 0.01, \*\*\**p* < 0.001).

## Acknowledgments

This work was supported by Research and Develop Program, West China Hospital of Stomatology Sichuan University (No. LCYJ2019-19, RD-03-202006); the National Natural Science Foundation of China (No. 81970965)

## Supplements

**Fig. S1.**
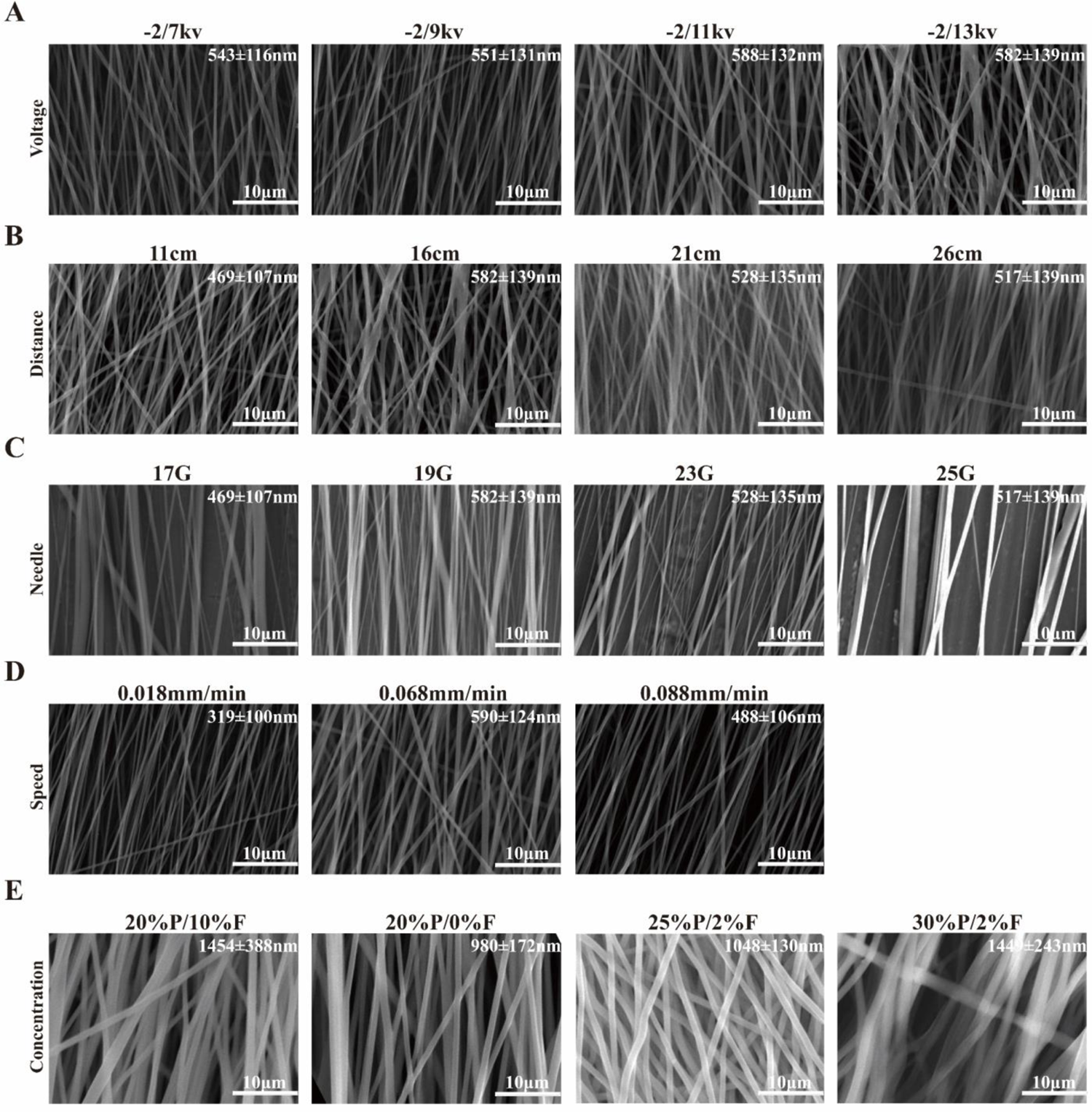
The diameter of the electrospinning fiber could be changed by adjusting (A) the voltage, (B) the distance from the tip to the collector, (C) the inner diameter of the needle, (D) the injection rate and (E) the concentration of the solution. P: PLGA, poly (lactide-co-glycolide); F: FC, fish collagen.

**Fig. S2.**
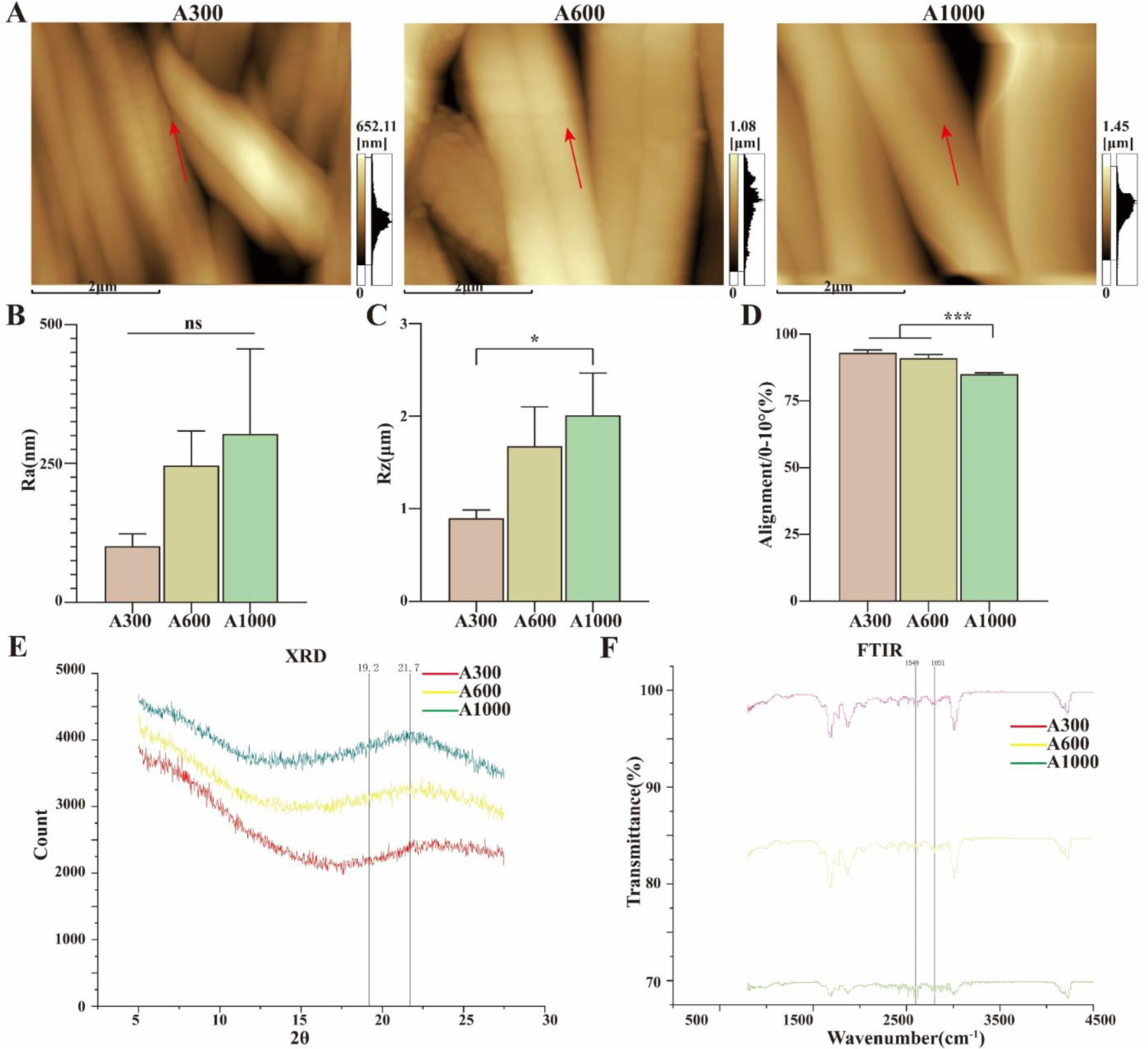
(A) AFM images and (B-C) corresponding roughness analysis of different diameter aligned electrospinning membranes; (D) Alignment analysis of various membranes; (E) XRD patterns of various membranes: the results showed there was no special peak, indicating that there was no formation of special crystal; (F) FTIR spectra of various membranes: the absorption of amide I and amide II in FC appeared in 1651 cm^-1^ and 1549 cm^-1^, respectively. Those absorption peaks in FC/PLGA (A300, A600 and A1000) have a weak shift, appearing at 1650 cm^−1^ and 1561 cm^−1^, respectively, indicating that the amino group in the FC most likely has hydrogen bonding with the PLGA molecular chain.

**Fig. S3.**
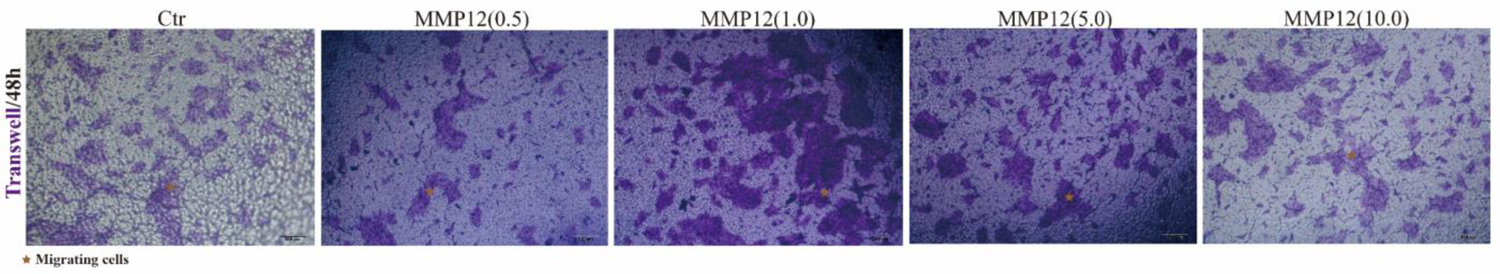
Screening of the optimal concentration of MMP12 for keratinocyte migration.

**Fig. S4.**
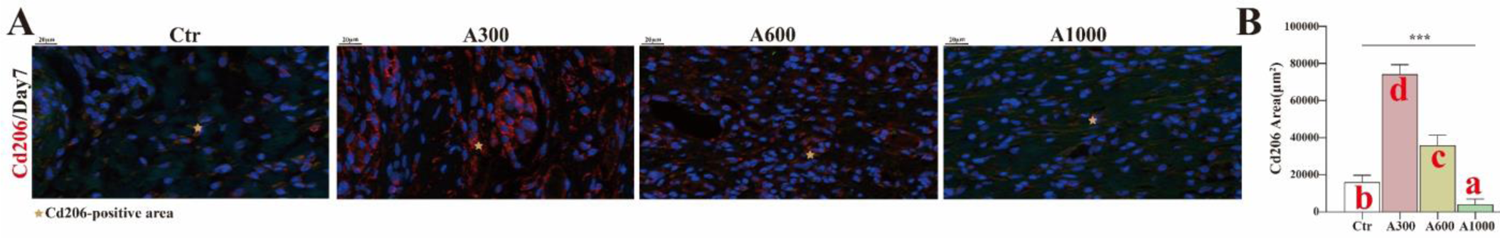
(A) Cd206-positive images of IF staining of rat skin defects and (B) corresponding analysis. Different letters indicate significant differences.

**Fig. S5.**
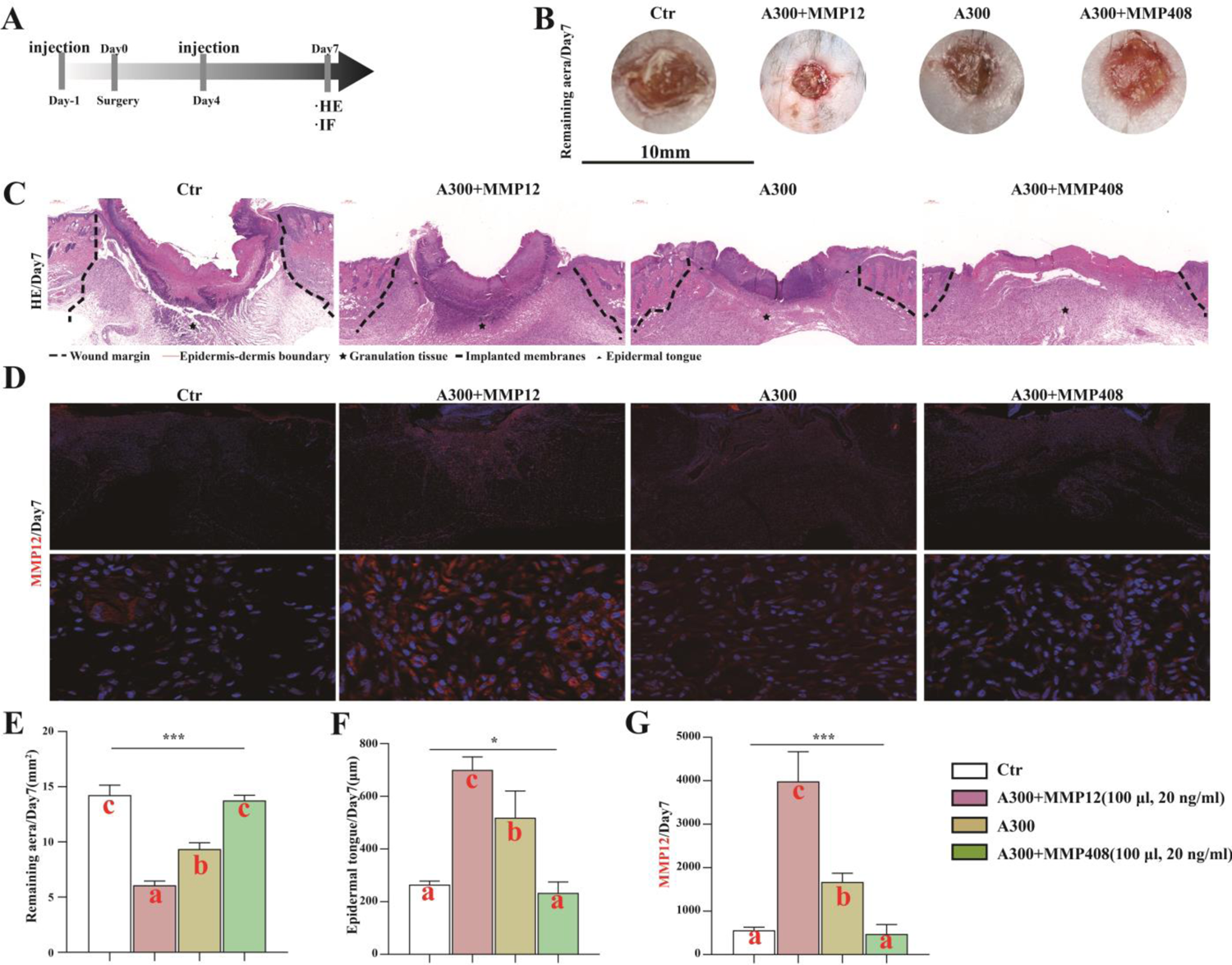
(A) Workflow for verifying the function of MMP12 in mice; (B) Macroscopic images of mouse skin defects and (E) corresponding analysis; (C) Residual gap images of H&E of mouse skin defects and (F) corresponding analysis;(D) MMP12-positive images of IF and (G) corresponding analysis. Different letters indicate significant differences.

**Table S1.** **Electrospinning parameters of Various Membranes**

| Diameter Group | Solutions | Voltage | Distance | Speed | Time |
| --- | --- | --- | --- | --- | --- |
| Small(A300) | PLGA (20% w/v) FC (2% w/v) | -2/5kv | 11cm | 0.018mm/min | 8h |
| Medium(A600) | PLGA (20% w/v) FC (2% w/v) |  | 16cm | 0.068mm/min | 4h |
| Large(A1000) | PLGA (25% w/v) FC (2% w/v) |  | 16cm | 0.068mm/min | 4h |

**Table S2.**
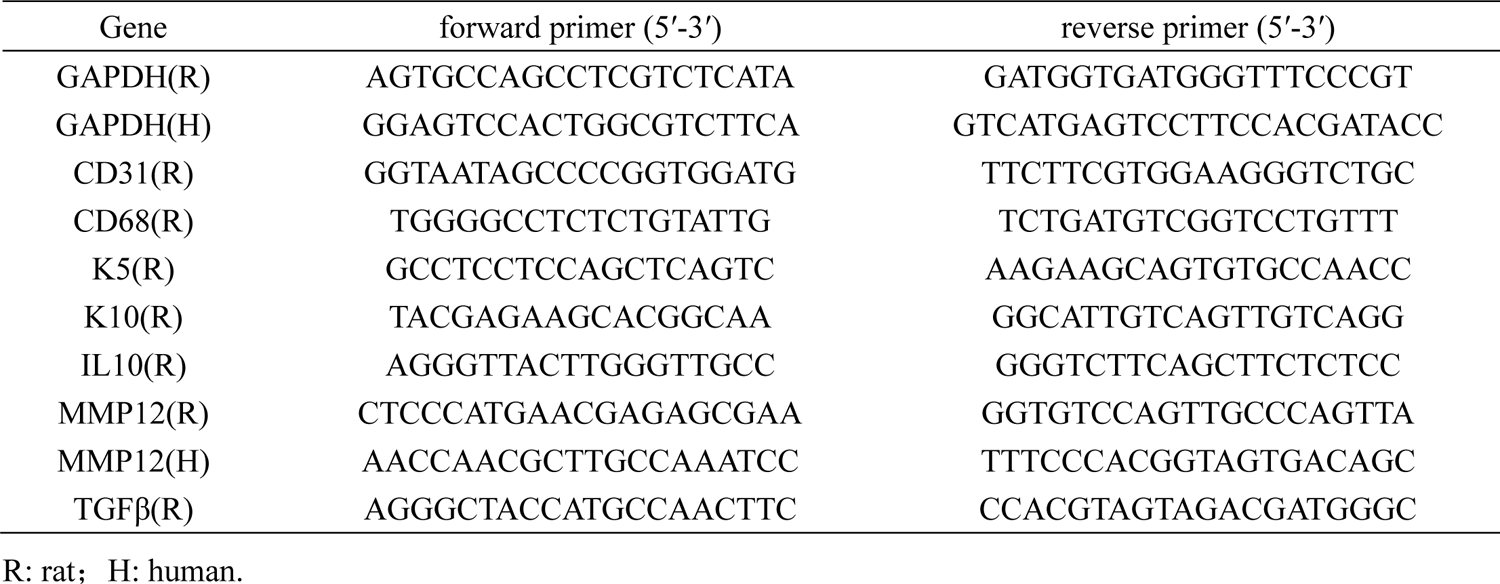
**Sequences of Primers Utilized in the Study**

## References

1. Amirsadeghi A, Jafari A, Eggermont L J, et al. Vascularization strategies for skin tissue engineering[J]. Biomater Sci, 2020, 8(15): 4073–4094.

2. Zhao Y, Li Z, Song S, et al. Skin-Inspired Antibacterial Conductive Hydrogels for Epidermal Sensors and Diabetic Foot Wound Dressings[J]. Advanced Functional Materials, 2019, 29.

3. Sen C K, Gordillo G M, Roy S, et al. Human skin wounds: a major and snowballing threat to public health and the economy[J]. Wound Repair Regen, 2009, 17(6): 763–71.

4. Blacklow S O, Li J, Freedman B R, et al. Bioinspired mechanically active adhesive dressings to accelerate wound closure[J]. Sci Adv, 2019, 5(7): eaaw3963.

5. Mascharak S, Desjardins-Park H E, Davitt M F, et al. Preventing Engrailed-1 activation in fibroblasts yields wound regeneration without scarring[J]. Science, 2021, 372(6540).

6. Savoji H, Godau B, Hassani M S, et al. Skin Tissue Substitutes and Biomaterial Risk Assessment and Testing[J]. Front Bioeng Biotechnol, 2018, 6: 86.

7. Wang P H, Huang B S, Horng H C, et al. Wound healing[J]. J Chin Med Assoc, 2018, 81(2): 94–101.

8. Cui L, Liang J, Liu H, et al. Nanomaterials for Angiogenesis in Skin Tissue Engineering[J]. Tissue Eng Part B Rev, 2020, 26(3): 203–216.

9. Larouche J, Sheoran S, Maruyama K, et al. Immune Regulation of Skin Wound Healing: Mechanisms and Novel Therapeutic Targets[J]. Adv Wound Care (New Rochelle), 2018, 7(7): 209–231.

10. Friedl P, Wolf K. Plasticity of cell migration: a multiscale tuning model[J]. J Cell Biol, 2010, 188(1): 11–9.

11. Scadden D T. Nice neighborhood: emerging concepts of the stem cell niche[J]. Cell, 2014, 157(1): 41–50.

12. Miguel S P, Figueira D R, Simões D, et al. Electrospun polymeric nanofibres as wound dressings: A review[J]. Colloids Surf B Biointerfaces, 2018, 169: 60–71.

13. Dellacherie M O, Seo B R, Mooney D J. Macroscale biomaterials strategies for local immunomodulation[J]. Nature Reviews Materials, 2019, 4(6): 379–397.

14. Sadtler K, Singh A, Wolf M T, et al. Design, clinical translation and immunological response of biomaterials in regenerative medicine[J]. Nature Reviews Materials, 2016, 1(7): 16040.

15. Juncos Bombin A D, Dunne N J, Mccarthy H O. Electrospinning of natural polymers for the production of nanofibres for wound healing applications[J]. Mater Sci Eng C Mater Biol Appl, 2020, 114: 110994.

16. Kishan A P, Cosgriff-Hernandez E M. Recent advancements in electrospinning design for tissue engineering applications: A review[J]. J Biomed Mater Res A, 2017, 105(10): 2892–2905.

17. Wang Y, Shi H, Qiao J, et al. Electrospun tubular scaffold with circumferentially aligned nanofibers for regulating smooth muscle cell growth[J]. ACS Appl Mater Interfaces, 2014, 6(4): 2958–62.

18. Nour S, Imani R, Chaudhry G R, et al. Skin wound healing assisted by angiogenic targeted tissue engineering: A comprehensive review of bioengineered approaches[J]. J Biomed Mater Res A, 2020.

19. Doyle A D, Carvajal N, Jin A, et al. Local 3D matrix microenvironment regulates cell migration through spatiotemporal dynamics of contractility-dependent adhesions[J]. Nat Commun, 2015, 6: 8720.

20. Liu Y, Franco A, Huang L, et al. Control of cell migration in two and three dimensions using substrate morphology[J]. Exp Cell Res, 2009, 315(15): 2544–57.

21. Xie J, Macewan M R, Ray W Z, et al. Radially aligned, electrospun nanofibers as dural substitutes for wound closure and tissue regeneration applications[J]. ACS Nano, 2010, 4(9): 5027–36.

22. Alford P W, Nesmith A P, Seywerd J N, et al. Vascular smooth muscle contractility depends on cell shape[J]. Integr Biol (Camb), 2011, 3(11): 1063–70.

23. Hu C, Chu C, Liu L, et al. Dissecting the microenvironment around biosynthetic scaffolds in murine skin wound healing[J]. Science Advances, 2021, 7(22): eabf0787.

24. Abrigo M, Mcarthur S L, Kingshott P. Electrospun nanofibers as dressings for chronic wound care: advances, challenges, and future prospects[J]. Macromol Biosci, 2014, 14(6): 772–92.

25. Mcbeath R, Pirone D M, Nelson C M, et al. Cell shape, cytoskeletal tension, and RhoA regulate stem cell lineage commitment[J]. Dev Cell, 2004, 6(4): 483–95.

26. Xie J, Shen H, Yuan G, et al. The effects of alignment and diameter of electrospun fibers on the cellular behaviors and osteogenesis of BMSCs[J]. Mater Sci Eng C Mater Biol Appl, 2021, 120: 111787.

27. Jafari A, Amirsadeghi A, Hassanajili S, et al. Bioactive antibacterial bilayer PCL/gelatin nanofibrous scaffold promotes full-thickness wound healing[J]. Int J Pharm, 2020, 583: 119413.

28. Kumbar S G, Nukavarapu S P, James R, et al. Electrospun poly(lactic acid-co-glycolic acid) scaffolds for skin tissue engineering[J]. Biomaterials, 2008, 29(30): 4100–7.

29. Liu Y, Ji Y, Ghosh K, et al. Effects of fiber orientation and diameter on the behavior of human dermal fibroblasts on electrospun PMMA scaffolds[J]. J Biomed Mater Res A, 2009, 90(4): 1092–106.

30. Hodgkinson T, Yuan X F, Bayat A. Electrospun silk fibroin fiber diameter influences in vitro dermal fibroblast behavior and promotes healing of ex vivo wound models[J]. J Tissue Eng, 2014, 5: 2041731414551661.

31. Saino E, Focarete M L, Gualandi C, et al. Effect of electrospun fiber diameter and alignment on macrophage activation and secretion of proinflammatory cytokines and chemokines[J]. Biomacromolecules, 2011, 12(5): 1900–11.

32. Zheng X, Xin L, Luo Y, et al. Near-Infrared-Triggered Dynamic Surface Topography for Sequential Modulation of Macrophage Phenotypes[J]. ACS Appl Mater Interfaces, 2019, 11(46): 43689–43697.

33. Chen H, Qian Y, Xia Y, et al. Enhanced Osteogenesis of ADSCs by the Synergistic Effect of Aligned Fibers Containing Collagen I[J]. ACS Appl Mater Interfaces, 2016, 8(43): 29289–29297.

34. Jin S, Sun F, Zou Q, et al. Fish Collagen and Hydroxyapatite Reinforced Poly(lactide-co-glycolide) Fibrous Membrane for Guided Bone Regeneration[J]. Biomacromolecules, 2019, 20(5): 2058–2067.

35. Gittens R A, Olivares-Navarrete R, Mclachlan T, et al. Differential responses of osteoblast lineage cells to nanotopographically-modified, microroughened titanium-aluminum-vanadium alloy surfaces[J]. Biomaterials, 2012, 33(35): 8986–94.

36. Chen Z, Bachhuka A, Han S, et al. Tuning Chemistry and Topography of Nanoengineered Surfaces to Manipulate Immune Response for Bone Regeneration Applications[J]. ACS Nano, 2017, 11(5): 4494–4506.

37. Mulholland E J. Electrospun Biomaterials in the Treatment and Prevention of Scars in Skin Wound Healing[J]. Front Bioeng Biotechnol, 2020, 8: 481.

38. Stachewicz U, Bailey R, Wang W, et al. Size dependent mechanical properties of electrospun polymer fibers from a composite structure[J]. Polymer, 2012, 53: 5132.

39. Tan E P, Ng S Y, Lim C T. Tensile testing of a single ultrafine polymeric fiber[J]. Biomaterials, 2005, 26(13): 1453–6.

40. Giljean S, Bigerelle M, Anselme K. Roughness statistical influence on cell adhesion using profilometry and multiscale analysis[J]. Scanning, 2014, 36(1): 2–10.

41. Miguel S P, Sequeira R S, Moreira A F, et al. An overview of electrospun membranes loaded with bioactive molecules for improving the wound healing process[J]. Eur J Pharm Biopharm, 2019, 139: 1–22.

42. Zhong S, Teo W E, Zhu X, et al. An aligned nanofibrous collagen scaffold by electrospinning and its effects on in vitro fibroblast culture[J]. J Biomed Mater Res A, 2006, 79(3): 456–63.

43. Chomachayi M D, Solouk A, Mirzadeh H. Electrospun silk-based nanofibrous scaffolds: fiber diameter and oxygen transfer[J]. Prog Biomater, 2016, 5: 71–80.

44. Tian F, Hosseinkhani H, Hosseinkhani M, et al. Quantitative analysis of cell adhesion on aligned micro- and nanofibers[J]. J Biomed Mater Res A, 2008, 84(2): 291–9.

45. Lavenus S, Pilet P, Guicheux J, et al. Behaviour of mesenchymal stem cells, fibroblasts and osteoblasts on smooth surfaces[J]. Acta Biomater, 2011, 7(4): 1525–34.

46. Baker B M, Trappmann B, Wang W Y, et al. Cell-mediated fibre recruitment drives extracellular matrix mechanosensing in engineered fibrillar microenvironments[J]. Nat Mater, 2015, 14(12): 1262–8.

47. Franz S, Rammelt S, Scharnweber D, et al. Immune responses to implants - a review of the implications for the design of immunomodulatory biomaterials[J]. Biomaterials, 2011, 32(28): 6692–709.

48. Anderson J M, Rodriguez A, Chang D T. Foreign body reaction to biomaterials[J]. Semin Immunol, 2008, 20(2): 86–100.

49. Chu C, Liu L, Rung S, et al. Modulation of foreign body reaction and macrophage phenotypes concerning microenvironment[J]. J Biomed Mater Res A, 2019.

50. Bryers J D, Giachelli C M, Ratner B D. Engineering biomaterials to integrate and heal: the biocompatibility paradigm shifts[J]. Biotechnol Bioeng, 2012, 109(8): 1898–911.

51. Sahana T G, Rekha P D. Biopolymers: Applications in wound healing and skin tissue engineering[J]. Mol Biol Rep, 2018, 45(6): 2857–2867.

52. Babakhani A, Nobakht M, Pazoki Torodi H, et al. Effects of Hair Follicle Stem Cells on Partial-Thickness Burn Wound Healing and Tensile Strength[J]. Iran Biomed J, 2020, 24(2): 99–109.

53. Chen F, Zhang Q, Wu P, et al. Green fabrication of seedbed-like Flammulina velutipes polysaccharides-derived scaffolds accelerating full-thickness skin wound healing accompanied by hair follicle regeneration[J]. Int J Biol Macromol, 2021, 167: 117–129.

54. Huang S M, Wu C S, Chiu M H, et al. High glucose environment induces M1 macrophage polarization that impairs keratinocyte migration via TNF-α: An important mechanism to delay the diabetic wound healing[J]. J Dermatol Sci, 2019, 96(3): 159–167.

55. Wolf M, Maltseva I, Clay S M, et al. Effects of MMP12 on cell motility and inflammation during corneal epithelial repair[J]. Exp Eye Res, 2017, 160: 11–20.

56. Chan M F, Li J, Bertrand A, et al. Protective effects of matrix metalloproteinase-12 following corneal injury[J]. J Cell Sci, 2013, 126(Pt 17): 3948–60.

57. Shapiro S D, Kobayashi D K, Ley T J. Cloning and characterization of a unique elastolytic metalloproteinase produced by human alveolar macrophages[J]. J Biol Chem, 1993, 268(32): 23824–9.

58. Daley J M, Brancato S K, Thomay A A, et al. The phenotype of murine wound macrophages[J]. J Leukoc Biol, 2010, 87(1): 59–67.

59. Elliott M R, Koster K M, Murphy P S. Efferocytosis Signaling in the Regulation of Macrophage Inflammatory Responses[J]. J Immunol, 2017, 198(4): 1387–1394.

60. Usui M L, Mansbridge J N, Carter W G, et al. Keratinocyte migration, proliferation, and differentiation in chronic ulcers from patients with diabetes and normal wounds[J]. J Histochem Cytochem, 2008, 56(7): 687–96.

61. Liu S-D, Li D-S, Yang Y, et al. Fabrication, mechanical properties and failure mechanism of random and aligned nanofiber membrane with different parameters[J]. Nanotechnology Reviews, 2019, 8: 218–226.

62. Da Cunha M R, Menezes F A, Dos Santos G R, et al. Hydroxyapatite and a New Fibrin Sealant Derived from Snake Venom as Scaffold to Treatment of Cranial Defects in Rats[J]. Materials Research-Ibero-American Journal of Materials, 2015, 18(1): 196–203.

